# Drosophila glue protects from predation

**DOI:** 10.1101/2020.12.23.424162

**Authors:** Flora Borne, Stéphane R. Prigent, Mathieu Molet, Virginie Courtier-Orgogozo

## Abstract

Animals can be permanently attached to a substrate in terrestrial environments at certain stages of their development. Pupa adhesion has evolved multiple times in insects and is thought to maintain the animal in a place where it is not detectable by predators. Here, we investigate whether pupa adhesion in Drosophila can also protect the animal by preventing potential predators from detaching the pupa. We measured the adhesion of Drosophila species sampled from the same area and found that pupa adhesion varies among species, which can be explained by different glue production strategies. Then, we compared attached and manually detached pupae in both field and laboratory assays to investigate the role of pupa adhesion to prevent predation. First, we found that attached pupae remain on site 30 % more than detached pupae in the field after three days, probably because they are less predated. Second, we observed that attached pupae are less efficiently predated by ants in the laboratory: they are not carried back to the ant nest and more ants are needed to consume them onsite. Our results show that pupa adhesion can prevent the animal from being taken away by predators and is crucial for Drosophila fly survival.

## Background

Multiple animals are immobile and permanently attached to a substrate. Most of them are found in aquatic environments (1) and within living hosts for parasites. In terrestrial environments, animals can also be attached to a substrate during certain stages of development where no feeding from outside nutrients is required. Eggs from multiple invertebrate species are fixed on leaves, wood or on the host tissue in some parasitic species (2–4). This is also the case of pupae, a non-feeding life stage of holometabolous insects between the larval and adult stages (5).

Modes of attachment and the ability to stick to the substrate appear to change rapidly during animal evolution. In stick and leaf insects gluing eggs to a substrate has evolved independently seven times (6). Wasp parasitoids have evolved different pupation strategies consisting in hanging their cocoon to a leaf (*Meteorus pulchricomis*) or attaching their cocoon on a leaf (*Microplitis sp*) (7). In flies, pupae of certain species such as *Drosophila melanogaster* and *Phormia regina* are glued to a substrate whereas others such as *Musca domestica*, *Calliphora erythrocephala* or *Sarcophaga falculata* are not (8).

Permanent attachment of eggs and pupae has been associated with several functions. First, attachment can allow the organisms to remain in a favorable environment. Females of many butterfly species choose to lay and attach their eggs directly on the host plant on which their larvae will start feeding (3,9). Butterflies laying their eggs during winter have evolved different strategies to avoid their eggs to be blown away if the dead host leaf falls far away from the plant. They lay their eggs on herbal or wooden substrates near the host plant or they glue their eggs less strongly to the host leaf so that the eggs would detach from the dead leaf and fall close to the host plant in case of strong wind (2,9,10). Second, hanging chrysalis may facilitate adult emergence (11). Third, attachment may protect immobile animals from predation in various ways. Permanent attachment, when associated with clumping behaviors, in which individuals of a particular species group closely to one another, can confer a better protection from predators. For example, in a freshwater caddisfly, pupal grouping behavior with conspecifics confers protection against a planarian flatworm predator (12). Attachment can also prevent predators from accessing the immobile animal. Cocoons of the parasitoid *Meteorus pulchricornis* are less predated when they are hanging than when they are artificially attached to leaves (14). To our knowledge, nothing is known about the function of pupal attachment in Diptera. In this study, we investigated whether the glue attaching Drosophila pupae can protect them from predation.

Drosophila larvae produce a glue right before pupariation which allows the pupa to stay attached to a substrate during metamorphosis. After expectoration, the glue spreads between the body and the substrate and dries within a few minutes (15). This glue is made of a few proteins called salivary gland secreted proteins (Sgs) which have evolved rapidly across and within species (16–19).

In Drosophila, pupae are found on rotten fruits (20), on or below the soil surface (21–23) and even on beer glass bottles (24). Pupation sites are usually close to the ground, thus accessible to ground dwelling species. The most common predators of fruit fly pupae are ants, rove beetles and spiders (25–27). Birds and small mammals were also found to prey on fruit fly pupae (28,29). To our knowledge, all studies on pupa predation in Drosophila were performed on *D. suzukii* (21,23,30). In these analyses, ants and spiders were the most common predators of pupae and ground beetles, earwigs and crickets were identified as potential natural predators. As ants were previously observed to dig up and carry pupae out of the soil (23), we hypothesized that pupa adhesion to a substrate might have another, yet unexplored, effect against predation: preventing potential predators from taking the animal away. Here, we first compared the adhesion strength of pupae from different drosophila species from the same ecological community to explore whether different species have evolved different attachment strategies. Then, we compared the ability of attached and detached pupae of one of these species to remain on site in a natural environment. Finally, we compared the ability of attached and detached pupae of two of these species to resist predation in the laboratory using the most common natural predator found in the field, an ant species.

## Materials and Methods

### Fly culture

Flies were cultured at 25 °C in plastic vials on standard medium [4 liters: 83.5 g yeast, 335.0 g cornmeal, 40.0 g agar, 233.5 g saccharose, 67.0 mL Moldex, 6.0 ml propionic acid]. For *D. suzukii*, this medium was supplemented with 200 g of D-glucose.

### Fly collection

Drosophila flies were collected at the Bois de Vincennes in Paris, France. On July 3 2020, five traps made from 0.5-L plastic bottles were settled in the ornithological reserve (48°50’05.5”N; 2°26’11.4”E). Small holes were made in the sides of the bottles that allowed drosophilid flies to enter but prevented entry by larger insects. Traps were baited with pieces of banana and hung to tree branches or within the understorey vegetation. They were distantly placed, three of them in the forest, one at the margin of a meadow and another close to a pond (Table S1). Flies in the bottles were collected two days and five days later. On July 16 2020, drosophilid flies were also collected with a sweeping net over a compost at the forest services facilities. Collected flies were transferred with an aspirator into plastic vials containing a piece of humid tissue paper for the time of transportation. In the laboratory, flies were checked under a binocular stereomicroscope and isolated by species in culture vials with standard cornmeal. When species could not be precisely determined, females were isolated in small culture vials with instant Drosophila medium (Formula 4-24, Carolina Biological Supply Company, Burlington, NC, USA) and species were then identified based on key morphological characters in the male progeny. Results of the fly collections are presented in Table S1. Combining isofemale lines of the same species when necessary, we managed to obtain mass culture for most species, including *D. hydei*, *D. simulans* and *D. suzukii*. These three stocks were raised in the laboratory for 3-4 months at 25°C before being used in the experiments described below.

### Adhesion assays

Third instar wandering larvae were transferred on glass slides (Menzel Superfrost microscope glass slide, ThermoScientific™ #AGAB000080) with soft forceps and kept in a box with wet cotton. 15 to 21 h after transfer, pupae naturally attached to the glass slides with their ventral part adhering to the glass slide were used for the adhesion tests. The pull-off force necessary to detach the pupa from the glass slide was measured using a universal test machine (LS1S/ H/230V Lloyd Instruments) with a 5N force sensor (YLC-0005-A1 Lloyd Instruments), in a set up similar to the one published earlier (31). Double-sided adhesive tape (tesa, extra strong, #05681-00018) was attached to a cylindrical metal part in contact with the force sensor. The force was set to 0 before each run. The force sensor was moved down with a constant speed of 1 mm/min until it pressed the pupa with a force of 0.07 N (0.25 N for *D. hydei*) then let still at a force of 0.03 N (0.21 N for *D. hydei*) for 10s and finally moved up with a constant speed of 0.2 mm/s until the pupa was detached. Force-by-time curves were recorded using NEXYGENPlus software (Lloyd Instruments). We used the maximal force reached during the experiment, corresponding to the force at which the pupa was detached, as the adhesion force of the individual. Pupae whose pupal case broke during the assay (*D. suzukii*: 1/27, *D. simulans*: 2/37; *D. hydei*: 7/50) or pupae which were not detached (*D. suzukii*: 0/27, *D. simulans*: 5/37, *D. hydei*: 12/50) were excluded from the analysis. After pupa detachment, images of glue prints remaining on glass substrates were taken with a Keyence VHX-2000 Z20 x20 or x100. Contours of prints areas were digitized manually by the same person using imageJ (1.50d, java 1.8.0_212, 64-bit) (32). Pictures were anonymized for manual contour acquisition so that the digitizer did not know the genotype. We measured the surface of the print corresponding to the pupa-substrate interface as defined previously (31). Three prints for *D. suzukii* and one print for *D. simulans* were not detectable on the slides and were not used in the analysis.

To assess pupal size, we used pupae that were raised in the same condition as for the adhesion tests but that were not used for the tests. We imaged attached pupae from the dorsal view and measured the area of the pupal case as described to measure the glue area.

### Predation assay in the field

In the laboratory, *D. simulans* third instar wandering larvae were collected with soft forceps and let to pupate in Petri dishes (55-mm diameter) in a closed plastic box containing wet paper for 17 to 24 hr at room temperature. 15 larvae were placed in each dish. All Petri dishes were then brought to the field. For the condition ‘attached pupae’, a few pupae were removed in order to have exactly 10 pupae glued in the lid of the Petri dish. For the condition ‘detached pupae’, pupae were all detached from the dish with featherweight entomology forceps and exactly 10 detached pupae were kept in the lid. To distinguish between the two conditions (“non collées” versus “collées” in French), the letters ‘NC’ or ‘C’ were written on pieces of paper that were taped on the external surface of the lids to facilitate visualisation and counting of the pupae. The lids of the Petri dishes were put in the center of buckets previously installed in the ornithological reserve of Bois de Vincennes (Fig. S1). These 40 × 35 cm buckets contain local soil, leaf litter and vegetation; they are pierced at the bottom for water draining and are semi-buried (10 cm deep) (33). In total, 28 buckets were used and contained both one dish with ‘attached pupae’ and one dish with ‘detached pupae’ (56 Petri dishes in total). We randomly alternated the West / East orientation of the two conditions inside the buckets. Pupae were counted at 0h (September 8 2020, day 1 morning, at 11 am), 6h30 (day 1 afternoon), 22h30 (day 2 morning), 31h (day 2 afternoon), 47h (day 3 morning), and 54h (day 3 afternoon) after the start of the experiment without being touched. Animals present in the dishes at counting times were photographed and later identified based on the pictures (Table S3). Insects present in the dishes at the end of recording (54h) were collected and kept in 90% ethanol for identification.

### Ant predation assay in the laboratory

Seven colonies of *T. nylanderi* were collected on September 17 2020 in Bois de Vincennes (48°50’20.0”N 2°26’57.2”E), brought to the lab and allowed to move into artificial nests consisting of two microscope glass slides separated by a 1-mm auto-adhesive plastic foam harboring three chambers, covered with a black plastic film to maintain darkness. Each artificial nest was placed in a foraging area consisting of a plastic box (11.5 × 11.5 × 5.5 cm) as described in (33). Water was always available in a tube plugged with cotton. Colonies were fed frozen Drosophila and diluted honey and then they were starved for 10 days until the beginning of the experiment, on October 13 2020. Prior and during the experiment, colonies were kept at room temperature on the bench and under indirect sunlight. Each day at around 9h30 am, and for 6 days (between October 13 and October 21 2020), one glass slide presenting two pupae was put into the foraging area of each colony, at about 10 cm from the entrances of the artificial nest. On each day, half of the colonies were given one slide with 2 *D. simulans* pupae and the other half one slide with 2 *D. suzukii* pupae, and we alternated species every day. We used the same two Drosophila lines as for our adhesion assays described above.

The glass slides were prepared as follows. Six third instar wandering larvae were transferred on glass slides (Menzel Superfrost microscope glass slide, ThermoScientific™ #AGAB000080) placed in a Petri dish kept in a box containing wet cotton and left to pupate for 14 to 19 hours. On each slide, only one pupa was kept attached, the other ones were detached slightly with soft forceps and one detached pupa was left on the slide. Pupae were about 1 cm apart from each other on the slide. Left / right positions of the two pupae were randomly assigned per colony and changed every day. The initial locations of the pupae were identified by a mark under the slide.

Ant foraging areas were checked by eye every 5 minutes for 3 hours after the glass slide with the 2 pupae was added. The number of ants in contact with the pupae were counted. If the pupa was brought to the nest, we noted the time when the pupa was present inside the nest for the first time. If the pupa was not brought to the nest, we noted the time when the pupa was fully consumed by the ants (no Drosophila body remnants visible by eye).

### Quantification and Statistical Analysis

We used R v3.6.1 (R Core Team 2015) to conduct our statistical analyses. Adhesion forces were not normally distributed and statistical differences in forces between species were tested by Kruskal-Wallis tests followed by multiple pairwise Wilcoxon tests. To test whether adhesion forces were correlated to pupa - substrate contact areas for each species, we performed standardized major axis regressions using *sma* function from the rsmatr-package in R (34). Slopes were compared among species. For predation assays in Bois de Vincennes and in the laboratory, means were compared using Wilcoxon tests because data were not normally distributed. In order to avoid a potential bucket effect on pupa disappearance in the predation assay in the field, we paired both “attached” and “detached” pupae conditions in each bucket and we subsequently used paired tests. As the number of detached pupae decreases towards the end of the experiment, predators might be more likely to prey upon attached pupae at later time points. Therefore we did not use a Cox proportional hazards model for survival analysis as the ratio of the hazard functions for individuals within the same bucket may change overtime. We did not correct P-values for multiple testing, as suggested by Nakagawa (2004).

## Results

### Glue adhesion strength varies between species from a same location

We collected drosophilid flies in the forest near Paris in June 2020 and noted the presence of 9 *Drosophila* species (Table S1). We established fly stocks of the most common species. Three of them, *Drosophila simulans*, *D. hydei* and *D. suzukii*, were assayed for pupal adhesion. We found that *D. simulans* detached at a median strength of 234.2 mN (Fig. 1A), similar to what has been found previously for its sister species *D. melanogaster* (31). We measured a lower adhesion for *D. suzukii* pupae with a median strength of 78.7 mN and higher adhesion for *D. hydei* with a median strength of 482.6 mN. Adhesion strength was significatively different between the three species (Kruskal-Wallis X^2^ = 63.77, *df* = 2, *p* = 1.4e-14, followed by all pairwise comparison Wilcoxon test, *p* < 0.001).

**Fig 1.**
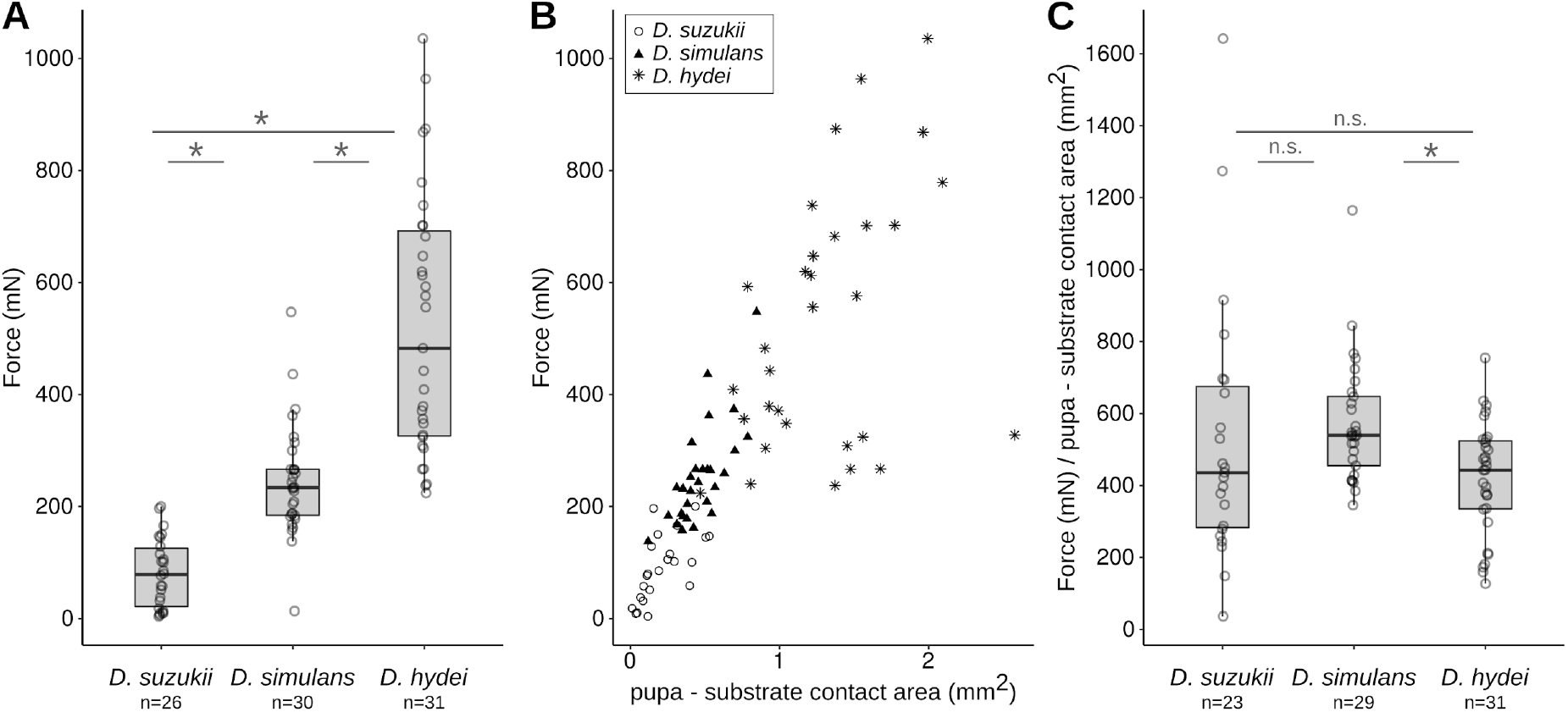
Pupa adhesion varies between species originating from the same location. (A) Adhesion strength of three Drosophila lines collected in Vincennes. Force indicates the force required to detach a pupa naturally attached to a glass slide. Each dot corresponds to a single pupa and n indicates the total number of pupae tested for each species. Ends of the boxes define the first and third quartiles. The black horizontal line represents the median. The vertical line on the top of the box extends to the largest value no further than 1.5 * IQR from the upper hinge of the box. The vertical line on the bottom of the box extends to the smallest value at most 1.5 * IQR of the hinge. (IQR: inter-quartile range is the distance between the first and the third quartiles). Data beyond the end of these lines are “outlying” points. * indicates significant differences between *D. suzukii* and *D. hydei*, *D. suzukii* and *D. simulans* and *D. simulans* and *D. hydei* (*p* < 0.05). (B) Relation between pupa adhesion force and pupa-substrate contact area. Each dot corresponds to a single pupa. *D. suzukii* pupae are represented as circles, *D. simulans* as triangles and *D. hydei* as stars. (C) Adhesion strength corrected by the pupa-substrate contact area. Boxplots and * as in 1A. n.s. indicates not significant (*p* > 0.05).

By examining the glue prints left by the pupae on glass slides after detachment, we found that adhesion forces correlated with the surface of the glue print delimiting the contact between the pupa and the glass slide for each species (Fig. 1B, *D. suzukii*: *R*^*2*^ = 0.41, *p* = 0.0009, *D. simulans*: *R*^2^ = 0.55, *p* = 5e-06, *D. hydei*, *R*^*2*^ = 0.17, *p* = 0.02). There was no difference in slope among the three species (*p* = 0.2) and the common slope was about 491, meaning that adhesion force increases by 491 mN for 1 mm^*2*^ increase of the pupa - substrate contact area for each species. After dividing the adhesion force by the surface of contact, we found a difference in adhesion between *D. simulans* and *D. hydei* (Fig. 1C, Kruskal-Wallis X^2^ = 9.61, *df* = 2, *p* = 0.008, followed by all pairwise comparison Wilcoxon test, *p* = 0.0008) but not between *D. simulans* and *D. suzukii* (*p* = 0.1) and *D. suzukii* and *D. hydei* (*p* = 0.6).

To test whether the production of the glue was related to the size of the pupa in each species, we imaged attached pupae raised in the same condition as the pupae used for the adhesion tests and measured the area of the pupal case. We found that the three species had different pupal size (Kruskal-Wallis X^2^ = 73.876, *df* = 2, *p* < 2e-16, followed by all pairwise comparison Wilcoxon test, *p* < 0.001) with *D. hydei* pupae presenting the biggest size (4.63 mm^2^), then *D. suzukii* (2.78 mm^2^) and *D. simulans* (2.12 mm^2^). The ratio of glue print area over pupal case area was 0.21 for *D. simulans*, 0.30 for *D. hydei* and 0.06 for *D. suzukii*, suggesting that *D. suzukii* pupae produce less glue relative to their size than *D. simulans* and *D. hydei*.

### Attached pupae are taken away less frequently than detached pupae in a semi-natural environment

To test whether glue attachment may protect pupae from predation in a semi-natural environment, we chose to use *D. simulans*, as we could obtain a large number of pupae from our fly strain. We compared the disappearance of pupae naturally attached to the plastic lid of Petri dishes with pupae mechanically detached from the lid. We placed two dishes containing respectively 10 attached and 10 detached pupae in 28 open buckets in Bois de Vincennes (two Petri dishes in each bucket) for 54 h and monitored the number of remaining pupae twice a day (Fig. 2A,B). At the end of the experiment, 10 pupae (median = 10) remained in the dish with attached pupae (all still attached) compared to 6-7 pupae (median = 6.5) in the dish with detached pupae. We found that attached pupae stayed significantly more in the Petri dishes than detached ones, with differences becoming significant from day 2 AM until the end of the experiment (Fig. 2F; paired Wilcoxon rank tests with continuity correction: day 1 PM: *V* = 9*, p* = 0.2; day 2 AM: *V* = 89.5, *p* = 0.02; day 2 PM: *V* = 92*, p* = 0.01; day 3 AM: *V* = 91*, p* = 0.002; day 3 PM: *V* = 110, *p* = 0.005). After three days, attached pupae remained on site 30 % more than detached pupae. During the countings, a few species were observed in the Petri dishes: *Temnothorax nylanderi* ants, red spider mites, cockroaches and woodlice (Table S5). Red spider mites were seen to fix themselves to both attached and detached pupae (Fig. 2C). *Temnothorax nylanderi* was the only species that was clearly seen consuming both attached and detached pupae in the dishes (Fig. 2D). Cockroaches were also observed, but only on the first day (Fig. 2E). Woodlice were also found in the dishes but never in contact with the pupae.

**Fig 2.**
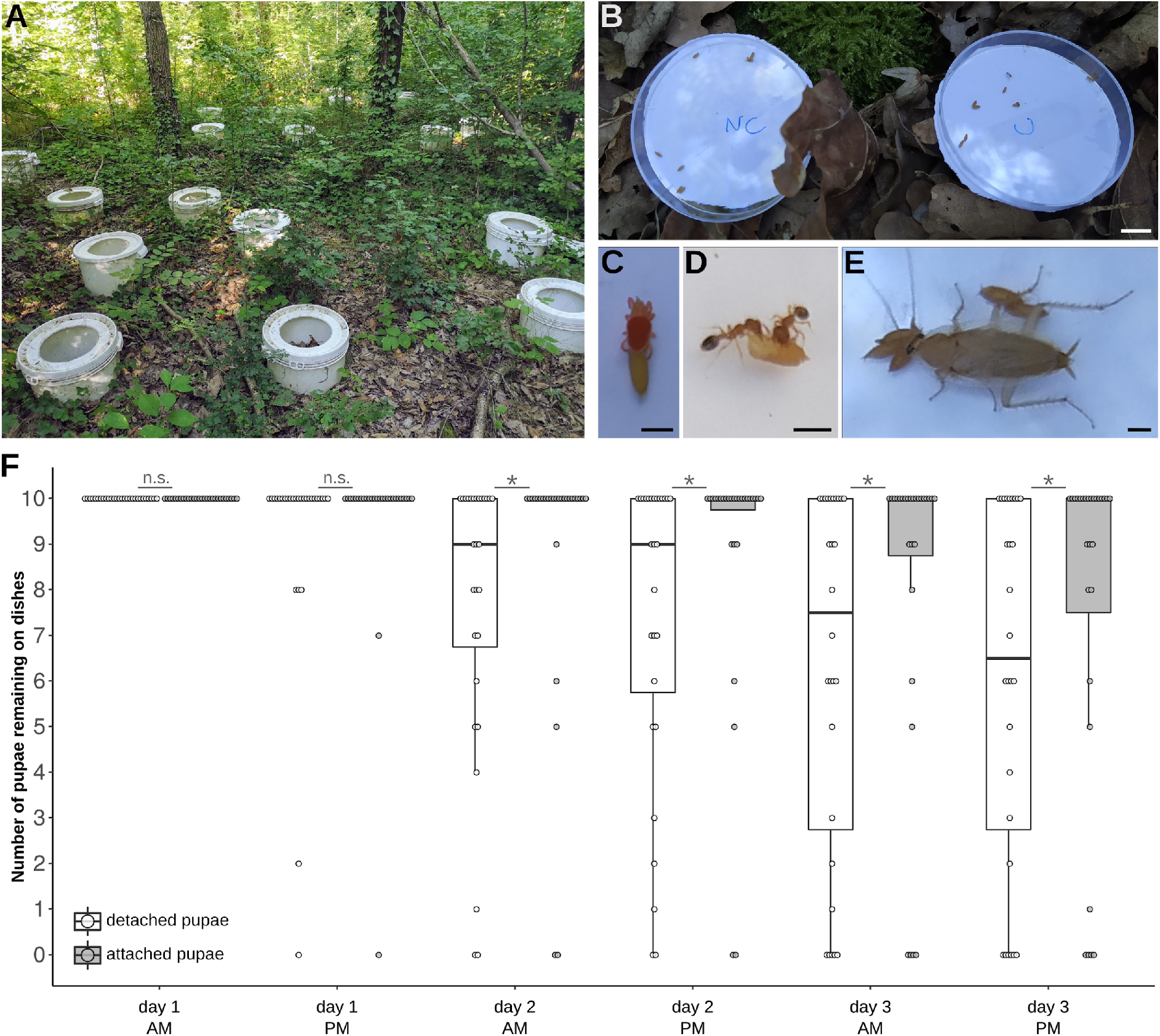
Attached pupae are taken away less frequently than detached pupae in nature. (A) Picture of the half-buried buckets installed in Bois de Vincennes. (B) Picture of two dishes placed in the center of a bucket and containing attached (“C”) or detached (“NC”) pupae. (C-E) Predators observed in the dishes during the experiments: a red mite spider (C), two *Temnothorax nylanderi* ants (D) and a cockroach (E). (F) Boxplot represents the number of pupae present in one dish at the counting time. Pupae were counted twice a day in the morning (AM) and in the afternoon (PM). Each dot represents the count for one dish. White boxes represent dishes with detached pupae and grey boxes dishes with attached pupae. Boxplots are defined as previously (Fig. 1A). * represents a significant difference between the number of remaining pupae between the attached and detached conditions (*p* < 0 .05, Wilcoxon test). Scale bars: (B) 1 cm, (C-E) 1 mm.

### Attached pupae are predated less efficiently by ants

To further understand how predators may act when they encounter an attached or a detached pupa, we decided to monitor in the laboratory pupae predation by the ant *Temnothorax nylanderi*, which was the most commonly found predator of *D. simulans* pupae in our field assay. Seven ant colonies were collected in Bois de Vincennes. After 10-day starvation, each ant colony was given on each day one glass slide with two pupae, an attached and a detached one. We used either two pupae of D. simulans (strongly attached) or two pupae of D. suzukii (loosely attached) and we alternated colonies each day. We examined the ant-Drosophila interactions every 5 minutes for 3 hours after adding the glass slide with pupae.

We found that in all replicates all pupae were consumed by the ants. For both fly species, we observed a difference between detached and attached pupae: detached pupae were mostly taken to the nest and eaten there while most of the attached pupae were eaten on site (Fig. 3A, number of pupae taken to the nest in *D. simulans:* detached 21/21, attached 3/21, *X*^*2*^ = 28.10, *df* = 1, *p* < 10^−6^; in *D. suzukii:* detached 19/21, attached 7/21, *X*^*2*^ = 12.22, *df* = 1, *p* = 0.0005). In 9 cases (6 for *D. suzukii* and 3 for *D. simulans*), both the attached and the detached pupa were taken to the nest; in all those cases the detached pupa was always taken to the nest first (about 37 min earlier for *D. suzukii* and 50 min earlier for *D. simulans* in median, Fig. 3A).

**Fig 3.**
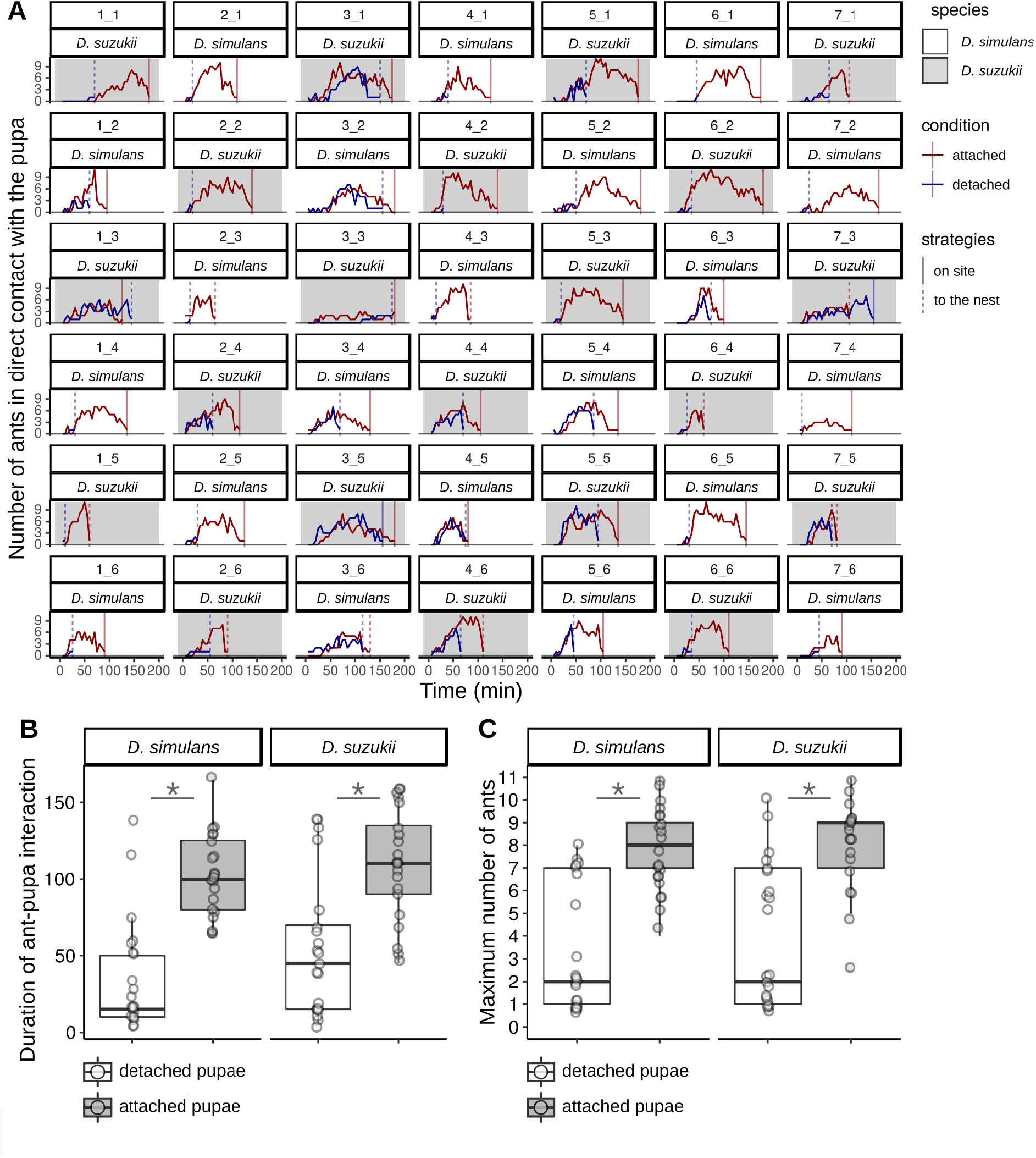
Attached pupae require more time and more ants to go away. (A) Number of ants in direct contact with the pupa over the duration of the experiment. Each cell represents one experiment with X_Y corresponding to the experiment with the colony X during the trial Y. Grey cells represent experiments using *D. suzukii* and white cells using *D. simulans*. Red lines represent the number of ants over time on attached pupa and blue lines on detached pupa. Vertical lines represent the time when the attached pupa (in red) or the detached pupa (in blue) arrives in the nest (dashed line) or is fully consumed outside the nest (full line) (B) Time during which ants are in contact with attached pupa (grey box) and detached pupa (white box) outside the nest (until the pupa is completely consumed outside the nest or enters into the nest). Each dot represents one experiment. (C) Maximum number of ants observed in contact with detached pupa (white box) and attached pupa (grey box) during the duration of the experiment. Each dot represents one experiment. Boxplots are defined as previously (Fig. 1A). * represents significant differences (*p* < 0.05, Wilcoxon tests).

Ants spent 15 min and 45 min (median) outside the nest in contact with *D. simulans* and *D. suzukii* detached pupae, respectively, while they spent respectively six and two times longer in contact with attached pupae (Fig. 3B, respectively 100 min and 110 min in *D. simulans* and *D. suzukii*, paired Wilcoxon rank tests with continuity correction *D. simulans: V* = 4.5, *p* = 0.0001; *D. suzukii: V* = 18, *p* = 0.0007). The maximum number of ants observed in direct contact with the fly pupa over the duration of the experiment was significantly higher for attached pupae than for detached pupae (Fig. 3A,C, median of 9 for *D. simulans* and 8 for *D. suzukii* for attached pupae compared to median of 2 for detached pupae in both species, paired Wilcoxon rank tests with continuity correction *D. simulans: V* = 2.5, *p* = 0.0002; *D. suzukii: V = 10.5, p* = 0.0007). The difference was still significant after correcting for the amount of time, by comparing the maximum number of ants in contact with the pupa until the first pupa is brought to the nest or fully consumed (paired Wilcoxon rank tests with continuity correction, *D. simulans*: 3 vs 2, *V* = 24, *p* = 0.02; *D. suzukii* 6 vs 2, *V* = 7, *p* = 0.004).

The time for the first ant to touch a pupa was slightly significantly different between attached and detached pupae only in *D. suzukii* (*D. suzukii*: 15 min vs 20 min, *V* = 31, *p* = 0.03; *D. simulans*: 15 min vs 20 min, *V* = 62.5, *p* = 0.3) but not different between *D. simulans* and *D. suzukii* (Wilcoxon rank tests with continuity correction, attached pupae: *W* = 195.5, *p* = 0.5; detached pupae: *W* = 200, *p* = 0.6). No differences were found between *D. simulans* and *D. suzukii* regarding the time until the detached pupae is brought to the nest from the start of the experiment (45 min vs. 65 min, *W* = 160.5, *p* = 0.296), the time that ants spent in contact with attached pupa (*W* = 194, *p* = 0.5) and detached pupa (*W* = 161, *p* = 0.1), the maximum number of ants over the duration of the experiment on detached pupae (*W* = 196.5, *p* = 0.5) or on attached pupae (*W* = 190.5, *p* = 0.5), the maximum number of ants until the first pupa disappeared on detached pupae (*W* = 231.5, *p* = 0.8) and the maximum number of ants on attached pupae (*W* = 265, *p* = 0.3).

## Discussion

### In the same environment, pupa adhesion strength varies among species

Using our previously published pull-force measurement assay (31), we provide here the first evidence that pupa adhesion varies between Drosophila species. Our result is in agreement with the rapid evolution of glue genes (17). Our analysis unravels at least two mechanisms leading to changes in the quantity of glue produced and resulting in changes in adhesion among species: *(i)* a change in body size (probably linked with a change in salivary gland size), as observed between *D. hydei* and *D. simulans*, *(ii)* a change in the amount of glue production independently of body size, as observed between *D. suzukii* and *D. simulans*. We note that our experiment might not reflect natural conditions as we have not tested adhesion on natural substrates and in natural habitat.

The variation we observed in adhesion force among individuals within a given species is much higher than measurement error (our universal test machine has an accuracy of ±0.5%) and could be due to individual variation in size, shape, weight, glue production, or position of the pupa relative to the substrate.

At the end of the larval stage, larvae stop feeding and start to search for a site to pupate. Pupation site choice during the larval stage is important for pupal survival. Pupation site preference has been thoroughly investigated in the lab (36–38) and more rarely in nature (39). This choice depends on abiotic factors such as temperature (37), darkness (40) or the nature of the substrates (41). In particular, *D. simulans* prefers to pupate on rough and humid surfaces while *D. hydei* prefers dry and smooth surfaces. *D. simulans* was often reported as pupating in fruits in the lab (36,38) and from field sampling (39) but other natural sites have not been investigated. In *D. suzukii*, recent studies in the field have found that pupae are present in the soil rather than in fruits (21,23). Additionally, pupation site preference depends on biotic factors and particularly on the presence of conspecifics and alien species. In *D. simulans* and *D. hydei* and in other Drosophila species, pupae are aggregated with conspecifics (39,42). In *D. simulans* and *D. buzzatii*, larvae change their site choice in presence of heterospecific larval cues (20). The differences we observed in adhesion strength among species may reflect differences in their ecology. For example, an adhesive substance might not be required for animals pupating on sticky substrates (rotten fruits and mushrooms, sap, …). Furthermore, species exhibiting distinct adhesion strengths in our laboratory conditions may nevertheless stick with similar forces in their respective natural habitats.

### Fixation of the pupa prevents predation

Comparing the disappearance of attached and detached *D. simulans* pupae in the field, we found that being attached allows pupae to stay on site more efficiently. Because pupae were contained within the lid of Petri dishes, we infer that they were not blown away by light wind, but we cannot be sure that missing pupae were predated. In our design, predators have the choice between attached and detached pupa within a bucket. This design reduces the possible effect of buckets when comparing attached and detached pupa disappearance.

The ant *Temnothorax nylanderi* was the main predator that we observed in the field consuming pupae. This observation is in agreement with previous studies which found that ants prey on fruit fly pupae (21,23,30). During our laboratory assay, *T. nylandei* ants predated attached and detached pupae with distinct behaviours: they brought most of the detached pupae to the nest while they ate the attached ones directly on site. The latter strategy requires ants to spend more time outside the nest and to recruit more foragers. The presence of parasites, predators and competitors in the wild would certainly make this strategy costly. Additionally, *T. nylanderi* is a solitary foraging species that recruits nestmates one by one (43). Under natural conditions, it would take a relatively long time to gather many foragers around the pupae. In our field assay, no more than two ants were observed together in a lid (Table S5). We found that ants act similarly on *D. suzukii* and *D. simulans* pupae, suggesting that there is no difference in strategy to predate loosely attached pupae such as *D. suzukii* or more strongly attached ones such as *D. simulans*, and that both species are equally attractive as preys.

To our knowledge, this study is the first to show that fly pupa adhesion can protect from predation. Our experiments are simple and can be easily applicable to other species, and not only for pupae but also for eggs, to check if this phenomenon is general in flies and insects.

### Alternative functions of Drosophila glue and alternative strategies to protect pupae from predation

Here, we demonstrated that pupal adhesion protects from predation by preventing predators like ants from taking the pupa away. Pupa adhesion may have alternative roles, such as maintaining the individual in a favorable environment (3, 9, 14) so that the pupa would be hidden from predators, protected from microorganism contaminants or/and have optimal conditions for pupal development. If not attached, the pupa could be moved away by abiotic factors such as wind or rain or biotic factors such as competitors. Pupa attachment could also help the adult to emerge from the pupal case, as suggested in butterflies (11) or facilitate pupal aggregation and thus dilution of predation risk as in freshwater caddisflies (12). Pupal congregation has been observed in Drosophila species (39,42) but its contribution to protection from predators has not been tested.

Pupal adhesion is only one of several strategies for pupae to escape predators. A common strategy is cryptic coloration to hide from visual predators (44,45). The brownish color of Drosophila pupae could contribute to hiding the animal when pupating in the leaf litter or in the soil. In some cases, pupae mimic non-living things such as leaves or sticks like the Common Maplet butterfly chrysalis (45). To avoid non visual predators, pupa has also evolved chemical defences either to chemically hide from predators (46) or to make the pupa toxic (47). Pupae have also evolved different types of physical defence (spin, hard pupal case, urticating hairs…). In Drosophila, the pupa is covered by a relatively thick (about 20 µm, (48)) and hard cuticle which has been hypothesized to protect the animal from predator attacks.

For the first time, we report that pupa adhesion varies among Drosophila species and that pupa attachment can protect from predation. Our results unravel a previously unknown important trait for Drosophila survival in the wild, the ability of pupae to firmly adhere to a substrate. Further studies of Drosophila glue combining genetic and phenotypic approaches should provide insight on the molecular basis for diverse bioadhesive properties, adapted to various habitats and climates.

## Acknowledgements

We are grateful to Michel Neff and the Division du Bois de Vincennes de la Ville de Paris for allowing us to use an area of wood land of the Réserve Ornithologique du Bois de Vincennes as an experimental plot. We thank Céline Bocquet for her help regarding collection and rearing of ant colonies, Manon Monier and Gauthier Beuque for their help with experiment installation in Vincennes, Aurélie Surtel for drawing the outlines of all the glue prints and Laurent Corté for discussions. The research leading to this paper has received funding from CNRS as part of the interdisciplinary action “Défi Adaptation du vivant à son environnement” 2020 and from the European Research Council under the European Community’s Seventh Framework Program (FP7/2007–2013 Grant Agreement no. 337579) to V.C.-O. F.B. was supported by a PhD fellowship from École normale supérieure Paris-Saclay. MM and the Vincennes setup were supported by the Institute of Ecology and Environmental Sciences - Paris and the Institut de la Transition environnementale de Sorbonne Université.

## Author Contributions

FB, MM and VCO designed the experiments, VCO supervised the project, SRP collected, identified and raised wild-caught flies, FB performed all the other experiments, FB, MM and VCO analyzed the results, FB wrote the original draft and all authors edited and contributed to the final version of this paper.

## Conflict of Interest Statement

The authors have no conflict of interest to declare.

## Data Availability Statement

Supplementary tables, raw data and scripts have been deposited in DRYAD (temporary link for private access: https://datadryad.org/stash/dataset/doi:10.5061/dryad.x3ffbg7hg). doi:10.5061/dryad.x3ffbg7hg

## Supplementary materials

**Figure S1.**
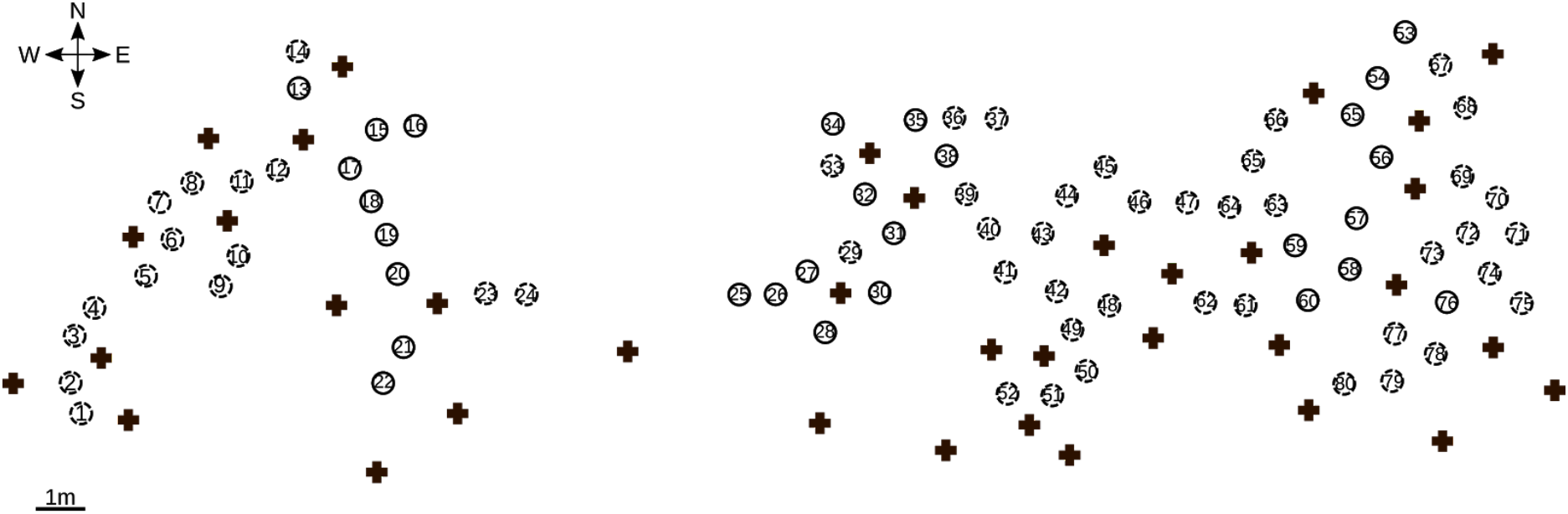
Distribution of the buckets in Bois de Vincennes. Each circle represents one bucket. Dashed lined circles indicate unused buckets and solid lined circles buckets used for the experiment. The number in the circle indicates the ID of the bucket. Crosses indicate the location of the trees. Orientation of the location is indicated in the top left corner.

**Table S1. List of Drosophila flies collected in Bois de Vincennes.** The number corresponds to the total number of individuals collected at each site. Traps were emptied twice, two days and five days after traps were set up in the field. GPS coordinates are shown for each trap and for the compost.

**Table S2. Adhesion force and pupa-substrate interface measurements.** Results of the adhesion assay performed on pupae from *D. suzukii*, *D. simulans* and *D. hydei* strain originating from Vincennes. Sample_ID column corresponds to a unique identification for each pupa, Temperature_assay, Humidity, Pressure_mba indicate respectively the room temperature (°C), the room humidity (%) and the atmospheric pressure (mba) at the moment of the assay. date_measurement and time_measurement correspond to the day and hours of the assay. date_substrate and time_substrate correspond to the day and hours at which larvae were put on the substrate. force_detachment_mN corresponds to the maximum force reached during the experiment in mN. Comment_on_this_sample reports particular observations: the pupa did not detach from the substrate (“not_detached”), the pupal case broke during the assay (“cuticle_broke”) or nothing special happened during the assay (“ok”). Area_px corresponds to the pupa-substrate contact surface measured in pixel and scale_px, scale_mm correspond respectively to the scale present on the picture in pixel and mm.

**Table S3. Pupal size measurement.** Measurements of the size of pupae of *D. simulans*, *D. suzukii*, *D. hydei*. ID column corresponds to a unique identification for each pupa. area corresponds to the area obtained by measuring the contour of the pupa in pixel. scale_px and scale_mm are defined as in Table S2.

**Table S4. Pupal count in the field experiment.** Results of the experiment performed in the field. bucket_ID column corresponds to a unique identification for each bucket. Orientation_C and Orientation_NC give, respectively, the orientation of attached and detached pupa within the bucket. Count_C and Count_NC indicate, respectively, the number of attached and detached pupa in a dish. Time indicates the time at which pupae were counted: t0 after 0h (8 September 2020, day 1 morning, at 11 am) t1 after 6h30 (day 1 PM), t2 after 22h30 (day 2 AM), t3 after 31h (day 2 PM), t4 after 47h (day 3 AM), and t5 after 54h (day 3 PM).

**Table S5. Animals observed in the dishes during the experiment with half-buried buckets in Bois de Vincennes**. The number indicates the number of dishes where the respective animals were observed. In total, 56 dishes were examined at each time point.

**Table S6. Ant count during predation assay in the laboratory.** The number indicates the number of ants observed in direct contact with the pupa at each time point. The column “X_Y” corresponds to the count over one experiment with the ant colony X at the trial Y.

**Table S7. Results table of the predation assay in the laboratory.** Each row corresponds to the description of the experiment for one pupa. condition column indicates the initial state of the pupa (“attached” or “detached”), strategies indicates whether the pupa was brought to the nest over the duration of the experiment (“to_nest”) or was never brought to the nest and consumed outside the nest (“on_site”). Time corresponds to the time at which the pupa was brought to the nest or fully consumed outside the nest in min. max_ant corresponds to the maximum number of ants observed in contact with the pupa over the duration of the experiment. orientation indicates whether the pupa was initially on the left or on the right side of the glass slide at the beginning of the experiment. first_ant corresponds to the time in min at which the first interaction between the pupa and an ant is observed. max_ant_at_first_done corresponds to the maximum number of ants observed in direct interaction with the pupa between the beginning and the time when the first pupa is brought to the nest or fully consumed outside the nest.

**R script. Drosophila_glue_predation.R** R script used to prepare the figures and run the statistical tests.

**pupa_prints** Folder containing pictures of the prints left by the pupae on glass slides after detachment during our adhesion assay force measurements.

**pupal_size** Folder containing the pictures of pupae used to measure the size of the pupae of the different species.

**Pictures_Vincennes** Folder containing pictures taken during and after the experiments in the field.

**Pictures_predation_lab** Folder containing pictures and video of the predation assays in the laboratory.

## Notes

https://datadryad.org/stash/share/SnN69p739EA9Q0KW-Tz6B7AZwEDkfRTZNIEedf6sZV8

### Competing Interest Statement

The authors have declared no competing interest.

### Summary of Updates

Text in Materials and Methods and Discussion updated. Fig. 2 updated. Fig. S1 (spatial distribution of the buckets) added.

## References

1. Wahl M. Living attached: aufwuchs, fouling, epibiosis. Fouling organisms of the Indian Ocean: biology and control technology. Oxf IBH Publ Co Put Ltd New Delhi. 1997 Jan 1;31–83.

2. Fordyce JA, Nice CC. Variation in butterfly egg adhesion: adaptation to local host plant senescence characteristics?: Variation in butterfly egg adhesion. Ecol Lett. 2002 Dec 13;6(1):23–7.

3. Hinton HE. Biology of insect eggs. 1st ed. Oxford [Eng.] ; New York: Pergamon Press; 1981. 3 p.

4. Voigt D, Gorb S. Egg attachment of the asparagus beetle Crioceris asparagi to the crystalline waxy surface of Asparagus officinalis. Proc R Soc B Biol Sci. 2010 Mar 22;277(1683):895–903.

5. Heming BS. Insect Development and Evolution [Internet]. Cornell University Press; 2003 [cited 2020 Dec 17]. 246–247 p. Available from: https://www.jstor.org/stable/10.7591/j.ctv75d5sv

6. Robertson JA, Bradler S, Whiting MF. Evolution of Oviposition Techniques in Stick and Leaf Insects (Phasmatodea). Front Ecol Evol. 2018 Dec 19;6:216.

7. Harvey JA, Gols R, Tanaka T. Differing Success of Defense Strategies in Two Parasitoid Wasps in Protecting Their Pupae Against a Secondary Hyperparasitoid. Ann Entomol Soc Am. 2011 Sep 1;104(5):1005–11.

8. Fraenkel G, Brookes VJ. The process by which the puparia of many species of flies become fixed to a substrate. Biol Bull. 1953 Dec;105(3):442–9.

9. Wiklund C. Egg-laying patterns in butterflies in relation to their phenology and the visual apparency and abundance of their host plants. Oecologia. 1984 Jul;63(1):23–9.

10. Hayes JL. A Study of the Relationships of Diapause Phenomena and Other Life History Characters in Temperate Butterflies. Am Nat. 1982 Aug;120(2):160–70.

11. Chapman RF. The Insects: structure and function. Cambridge (Mass.): Harvard University Press; 1982.

12. Wrona FJ, Dixon RWJ. Group Size and Predation Risk: A Field Analysis of Encounter and Dilution Effects. Am Nat. 1991;137(2):186–201.

13. Hieber CS. Spider cocoons and their suspension systems as barriers to generalist and specialist predators. Oecologia. 1992 Oct;91(4):530–5.

14. Shirai S, Maeto K. Suspending cocoons to evade ant predation in *Meteorus pulchricornis*, a braconid parasitoid of exposed-living lepidopteran larvae. Entomol Sci. 2009 Mar;12(1):107–9.

15. Beňová-Liszeková D, Beňo M, Farkaš R. Fine infrastructure of released and solidified *Drosophila* larval salivary secretory glue using SEM. Bioinspir Biomim. 2019 Jul 11;14(5):055002.

16. Beckendorf SK, Kafatos FC. Differentiation in the salivary glands of Drosophila melanogaster: Characterization of the glue proteins and their developmental appearance. Cell. 1976 Nov 1;9(3):365–73.

17. Da Lage J-L, Thomas GWC, Bonneau M, Courtier-Orgogozo V. Evolution of salivary glue genes in Drosophila species. BMC Evol Biol. 2019 Dec;19(1):36.

18. Korge G. Chromosome puff activity and protein synthesis in larval salivary glands of Drosophila melanogaster. Proc Natl Acad Sci. 1975 Nov 1;72(11):4550–4.

19. Korge G. Larval saliva in Drosophila melanogaster: Production, composition, and relationship to chromosome puffs. Dev Biol. 1977 Jul;58(2):339–55.

20. Beltramí M, Medina-Muñoz M, Pino F, Ferveur J-F, Godoy-Herrera R. Chemical Cues Influence Pupation Behavior of Drosophila simulans and Drosophila buzzatii in Nature and in the Laboratory. PloS One. 2012 Jun 21;7:e39393.

21. Ballman ES, Collins JA, Drummond FA. Pupation Behavior and Predation on Drosophila suzukii (Diptera: Drosophilidae) Pupae in Maine Wild Blueberry Fields. J Econ Entomol. 2017 Dec 5;110(6):2308–17.

22. Grossfield. Non-sexual behavior of Drosophila. In: The genetics and biology of Drosophila. Ashburner M, Wright TRF. London, New York, San Francisco: Academic Press; 1978. p. 3–126.

23. Woltz JM, Lee JC. Pupation behavior and larval and pupal biocontrol of Drosophila suzukii in the field. Biol Control. 2017 Jul;110:62–9.

24. Vouidibio J, Capy P, Defaye D, Pla E, Sandrin J, Csink A, et al. Short-range genetic structure of Drosophila melanogaster populations in an Afrotropical urban area and its significance. Proc Natl Acad Sci. 1989 Nov 1;86(21):8442–6.

25. Hennessey MK. Predation on wandering larvae and pupae of caribbean fruit fly (diptera: tephritidae) in guava and carambola grove soils. J Agric Urban Entomol. 1997;14(2):129–38.

26. Thomas DB. Predation on the soil inhabiting stages of the Mexican fruit fly. Southwest Entomol. 1995;20(1):61–71.

27. Urbaneja A, Marí FG, Tortosa D, Navarro C, Vanaclocha P, Bargues L, et al. Influence of Ground Predators on the Survival of the Mediterranean Fruit Fly Pupae, Ceratitis capitata, in Spanish Citrus Orchards. Biocontrol. 2006 Oct 3;51(5):611–26.

28. Bigler F, Neuenschwander P, Delucchi V, Michelakis S. Natural enemies of preimaginal stages of Dacus oleae Gmel.(Dipt., Tephritidae) in Western Crete. II. Impact on olive fly populations. Boll Lab Entomol Agrar Filippo Silvestri Italy. 1986;43:79–96.

29. Thomas DB. Survivorship of the Pupal Stages of the Mexican Fruit Fly Anastrepha ludens (Loew) (Diptera: Tephritidae) in an Agricultural and a Nonagricultural Situation. J Entomol Sci. 1993 Oct 1;28(4):350–62.

30. Gabarra R, Riudavets J, Rodríguez GA, Pujade-Villar J, Arnó J. Prospects for the biological control of Drosophila suzukii. BioControl. 2015 Jun;60(3):331–9.

31. Borne F, Kovalev A, Gorb S, Courtier-Orgogozo V. The glue produced by *Drosophila melanogaster* for pupa adhesion is universal. J Exp Biol. 2020 Apr 15;223(8):jeb220608.

32. Schneider CA, Rasband WS, Eliceiri KW. NIH Image to ImageJ: 25 years of image analysis. Nat Methods. 2012 Jul;9(7):671–5.

33. Honorio R, Doums C, Molet M. Manipulation of worker size diversity does not affect colony fitness under natural conditions in the ant Temnothorax nylanderi. Behav Ecol Sociobiol. 2020 Aug;74(8):104.

34. Warton DI, Duursma RA, Falster DS, Taskinen S. smatr 3–an R package for estimation and inference about allometric lines. Methods Ecol Evol. 2012;3(2):257–9.

35. Nakagawa S. A farewell to Bonferroni: the problems of low statistical power and publication bias. Behav Ecol. 2004 Nov 1;15(6):1044–5.

36. Erezyilmaz DF, Stern DL. Pupariation Site Preference Within and Between Drosophila Sibling Species. Evolution. 2013;67(9):2714–27.

37. Schnebel EM, Grossfield J. Temperature effects on pupation-height response in four Drosophila species group triads. J Insect Physiol. 1992 Oct 1;38(10):727–32.

38. Vandal NB, Siddalingamurthy GS, Shivanna N. Larval pupation site preference on fruit in different species of *Drosophila*. Entomol Res. 2008 Sep;38(3):188–94.

39. Beltramí M, Muñoz M, Arce D, Godoy-Herrera R. Drosophila pupation behavior in the wild. Evol Ecol. 2010 Mar 1;24.

40. Rizki MTM, Davis Charles G. Light as an Ecological Determinant of Interspecific Competition between Drosophila willistoni and Drosophila melanogaster. Am Nat. 1953 Nov 1;87(837):389–92.

41. Godoy-Herrera R, Luis S-C. The behavior of sympatric Chilean populations of Drosophila larvae during pupation. Genet Mol Biol. 1998 Mar 1;21.

42. Ringo J, Dowse H. Pupation Site Selection in Four Drosophilid Species: Aggregation and Contact. J Insect Behav. 2012 Nov 1;25.

43. Glaser S, Grueter C. Ants (Temnothorax nylanderi) adjust tandem running when food source distance exposes them to greater risks. Behav Ecol Sociobiol. 2018 Feb 19;72:40.

44. Gaitonde N, Joshi J, Kunte K. Evolution of ontogenic change in color defenses of swallowtail butterflies. Ecol Evol. 2018 Oct;8(19):9751–63.

45. Lindstedt C, Murphy L, Mappes J. Antipredator strategies of pupae: how to avoid predation in an immobile life stage? Philos Trans R Soc B Biol Sci. 2019 Oct 14;374(1783):20190069.

46. Mizuno T, Hagiwara Y, Akino T. Chemical tactic of facultative myrmecophilous lycaenid pupa to suppress ant aggression. Chemoecology. 2018 Dec;28(6):173–82.

47. Deyrup ST, Eckman LE, Lucadamo EE, McCarthy PH, Knapp JC, Smedley SR. Antipredator activity and endogenous biosynthesis of defensive secretion in larval and pupal Delphastus catalinae (Horn) (Coleoptera: Coccinellidae). Chemoecology. 2014 Aug;24(4):145–57.

48. Mitchell HK, Weber-Tracy UM, Schaar G. Aspects of cuticle formation in Drosophila melanogaster. J Exp Zool. 1971;176(4):429–43.

